# Gray matter Volume Modulates the Effect of Acute Physical Activity on Reading Comprehension and Cognitive Load in Adolescents –The Cogni-Action Project

**DOI:** 10.64898/2026.03.31.715252

**Authors:** Ricardo Martínez-Flores, Hans Supèr, Javier Sanchez-Martinez, Patricio Solis-Urra, Benjamin Tari, Romualdo Ibañez, Fabian Herold, Fred Paas, Myrto Mavilidi, Liye Zou, Carlos Cristi-Montero

## Abstract

**Background:** Physical activity has been associated with better reading comprehension and reduces cognitive load (CL), but the role of brain volume in modulating this relationship remains unclear. Therefore, this study aims to determine whether the gray matter volume in key regions modulates the effects of different physical activity modalities on reading comprehension and associated CL.

**Methods:** Thirteen male adolescents (12–13 years). Adolescents with MRI data participated in a randomized cross-over trial comparing three conditions: 1) sedentary behavior (SC, emulating a school class), 2) moderate-intensity continuous training (MICT), and 3) cooperative high-intensity interval training (C-HIIT), with physical activity conditions duration adjusted to match SC energy expenditure. Gray matter volumes were measured in the bilateral hippocampus, left pars opercularis, and the brainstem. CL was assessed via pupil dilation during reading using eye-tracking. Reading comprehension was measured through seven-question multiple-choice tests with expert-validated items.

**Results:** C-HIIT demonstrated superior effects on both CL and reading comprehension compared to MICT and SC, with significant brain volume modulation effects across all examined regions. Brain volume interactions with physical activity modalities systematically modified the pattern of cognitive responses, with C-HIIT consistently benefiting from these modulations, whereas the effects of MICT were generally attenuated.

**Conclusion:** This study suggests that selecting the appropriate physical activity modality may be relevant for cognitive outcomes during reading in adolescents. C-HIIT yielded lower CL and better reading comprehension, and these effects were not explained by brain volume alone but by its interaction with exercise modality.

## 1. Introduction

Appropriate cognitive development, including reading comprehension, during childhood and adolescence is critical for acquiring essential skills for success in later life (e.g., job success) and mental health (Eyre et al., 2024; John et al., 2022). Reading comprehension is a multifaceted cognitive process that involves not only basic decoding but also the integration of information into coherent mental representations, according to the construction–integration model, it unfolds as a structured process that increases in cognitive complexity (Follmer, 2018; Kintsch, 1988; Van Den Broek & Kendeou, 2024). Notably, accumulating evidence indicates a decline in reading performance among the global population of children and adolescents (OECD, 2023). For instance, the latest “Survey of Adult Skills – Reader’s Companion” by the Organization for Economic Co-operation and Development (OECD) highlights a persistent drop in foundational reading abilities (OECD, 2024). Similarly, results from the Programme for International Student Assessment (PISA) show that nearly 20% of 15-year-olds failed to reach the baseline reading proficiency needed for meaningful participation in industrialized societies (OECD, 2023). Further, poor reading skills during early life are also associated with a higher risk of cognitive decline and dementia later in life (John et al., 2022). Given those findings, it is important to support the optimal cognitive development early in life by appropriate means to improve key cognitive skills, such as reading comprehension, to foster broader societal development in line with the United Nations’ Sustainable Development Goals (Eyre et al., 2024).

Individual differences in brain structure, such as gray matter volume in the bilateral hippocampus, left pars opercularis, and brainstem, serve as robust predictors of reading comprehension, particularly during adolescence when these regions are maturing and susceptible to environmental influences like physical activity (Keller et al., 2024; Skoe et al., 2017). The brainstem is closely connected to classical biomarkers of text processing, such as pupil dilation, which serves as a reliable proxy for cognitive load (CL), thus larger pupil dilation indicates higher CL, whereas smaller dilation or constriction reflects lower CL (Joshi & Gold, 2020; Zou et al., 2023). Given the embryological link between the eye and brain, pupil size reflects neural activity during cognitive tasks of several parts of the brain, reinforcing the relevance of considering pupillometry in developmental cognitive (neuroscience) research especially given its non-invasive nature, minimal setup, and suitability for children and adolescents (Eckstein et al., 2017; Joshi & Gold, 2020; Moore et al., 2015; Zou et al., 2023).

The brainstem houses several key structures that regulate eye movements and pupil size, including the locus coeruleus, which is the brain’s primary source of norepinephrine thus is crucial for cognitive arousal (Khroud et al., 2024). The locus coeruleus modulates crucial cognitive functions related to reading, such as visual attention, through its widespread neural projections (Aston-Jones et al., 1999). Functionally, it is connected to the Edinger–Westphal nucleus, which mediates pupillary constriction by relaying signals to the iris sphincter muscle via the ciliary ganglion (Heiland Hogan et al., 2025). Moreover, the locus coeruleus receives excitatory input from the ventral hippocampus, which communicates with the medial prefrontal cortex to regulate attention and information processing (Tsetsenis et al., 2022; van den Brink et al., 2019). Thus, the pupil dilatations is a relatively easy-to-capture readout of the activation not only of specific brainstem regions but also other brain structures such as the hippocampus, and frontal and prefrontal areas, including to inferior frontal gyrus liked the pars opercularis, indicating that dilatation or constriction is related with increase or decrees CL respectably (Alnæs et al., 2014; Heiland Hogan et al., 2025; Tsetsenis et al., 2022).

Beyond its cognitive functions, the brainstem also plays a central role in physical activity. Although voluntary motor commands originate in the motor cortex, they are transmitted through brainstem circuits, making this region a crucial hub for movement control (Leiras et al., 2022). Notably, especially high-intensity physical activity has been shown to activate the locus coeruleus via its noradrenergic pathway, thereby may enhancing attentional control performance (Plini et al., 2023; Vazey et al., 2018). Previous evidence has shown that a single bout of physical activity can enhance brain function (Pontifex et al., 2019; Yu et al., 2021; Zou et al., 2023). However, its effects depend on the mode and intensity of the activity, emphasizing the importance of a dose-response relationship, as well as the need to investigate different modalities (e.g., continuous vs. intermittent) and intensities (e.g., moderate vs. vigorous) (Herold et al., 2019; Mavilidi et al., 2025). Furthermore, such physical activity has been associated with improved connectivity and responsiveness in several brain regions involved in CL and reading comprehension, including the hippocampus and the inferior frontal gyrus (Festa et al., 2023).

Building on this, a recent study found that the effects of physical activity on both reading comprehension and CL varied depending on the mode and intensity (i.e., continuous moderate-intensity vs. interval high-intensity physical activity) (Martínez-Flores et al., 2025). Specifically, interval high-intensity physical activity was associated with modulated pupillary responses, indicating lower CL during reading and better performance on comprehension questions targeting both surface code and situation model levels (Martínez-Flores et al., 2025). These findings suggest the recruitment of distinct neural systems involved in managing CL, and also point to the possibility that individual differences—such as the size of specific brain structures—may significantly contribute to variability in CL and reading comprehension outcomes (Broek et al., 2015).

In line with this, previous research has linked interindividual differences in brain anatomy, particularly in structures such as the inferior frontal gyrus and hippocampus (Keller et al., 2024; Skoe et al., 2017) with variations in reading comprehension and pupillary responses. This raises the possibility that such neuroanatomical factors could shape the cognitive benefits derived from acute physical activity. These observations align with a growing body of evidence, including longitudinal studies and meta-analysis, stressing the importance of considering individual differences in brain morphology (e.g., gray matter volume) when examining reading comprehension and the variables that influence it, especially during sensitive developmental stages such as childhood and adolescence (Eckert et al., 2016; Marks et al., 2024; Perdue et al., 2020).

Collectively, the this research seeks to deepen our understanding of how gray matter volume interact with physical activity to influence reading comprehension and the associated CL in adolescents. It addresses two major gaps in the current literature. First, although pupil dilation is a well-established biomarker of CL (van der Wel & van Steenbergen, 2018) little is known about how physical activity affects this relationship when individual brain structure is taken into account (Zou et al., 2023). In this sense, the vast majority of research explores the behavioral characteristics of physical activity with CL. However few studies connect physical activity and CL utilizing neurophysiological methods (i.e., pupil dilation). Second, most volumetric brain research has been conducted outside Latin America, leaving a significant geographical gap (Wassenaar et al., 2021). In this sense this study offers valuable insights on whether brain volume in specific regions modulates the effect of physical activity of varying intensity on CL and reading comprehension.

Therefore, this study examined whether gray matter volume in key regions modulates the effects of different physical activity modalities on reading comprehension and associated CL in early adolescents, a developmental stage of particular sensitivity for both brain maturation and the acquisition of academic skills. Four hypotheses were formulated. First, C-HIIT was expected to produce superior reading comprehension and lower CL than both MICT and the sedentary condition, consistent with evidence linking high-intensity interval exercise to greater noradrenergic activation and executive function benefits relevant to academic performance. Second, MICT was expected to outperform the sedentary condition on both outcomes, though to a lesser degree, reflecting the dose-dependent nature of the exercise-cognition relationship. Third, baseline gray matter volume in the examined regions was not expected to independently predict reading comprehension or CL, given that structural individual differences are thought to require appropriate physiological activation to manifest as functional advantages in learning contexts. Fourth, and centrally, gray matter volume was expected to significantly moderate the effect of physical activity modality on both outcomes, with the strongest modulation under C-HIIT, suggesting that greater structural capacity in reading-relevant regions may amplify the cognitive benefits that high-intensity exercise confers in school settings. Based on the functional specialization described in the construction-integration model and prior neuroimaging literature, region-specific modulation patterns were considered plausible: brainstem and pars opercularis volumes have been linked to foundational text processing, whereas hippocampal volume has been associated with higher-order comprehension and contextual integration (Keller et al., 2024; Kintsch, 1988). Because no prior study has directly examined the interaction between acute exercise and regional brain volume in this context, these region-level expectations were treated as exploratory rather than confirmatory.

## 2. Methods

This is a secondary analysis of data from the cross-sectional Cogni-Action Project, conducted between March 2017 and October 2019. A complete and extended version of the methodology is available in the Supplementary Material. Additionally, comprehensive methodological details have been previously published in two independent studies (Martínez-Flores et al., 2025; Solis-Urra et al., 2019). This study was registered in a clinical trial database and approved by the Ethics Committee of the authors’ institution. All phases were conducted in accordance with the Declaration of Helsinki.

### 2.1 Participants

Of the original sample (n = 1,296), two subsamples were recruited: 76 participants who underwent MRI neuroimaging (Solis-Urra et al., 2019) and 32 participants who completed an eye-tracking crossover trial (Martínez-Flores et al., 2025). The present study includes only participants for whom both MRI and eye-tracking data were available. Of the 32 crossover participants, 13 had usable MRI data, yielding a final sample of n = 13. This reflects data availability rather than a priori power calculations, as this is a secondary analysis combining two previously published datasets; original sample sizes and power justifications are reported in their respective studies.

For the MRI subsample, only right-handed adolescents were included to ensure homogeneity in hemispheric organization, given that handedness is associated with cortical asymmetries that could confound brain–behavior relationships (Solis-Urra et al., 2019). Participants were male students enrolled in 7th or 8th grade (12–13 years old) at a subsidized school of middle socioeconomic status, reflecting the feasibility constraints of the original crossover trial. Inclusion criteria required native Spanish speakers with normal or corrected-to-normal vision and no physical, psychiatric, or psychological disabilities, confirmed by physician evaluation. Full eligibility details are reported in Martínez-Flores et al. (2025).

Although the sample is small, the within-subjects crossover design provides 39 observations (13 × 3 conditions) and 9,556 datapoints, substantially reducing between-participant variance (Pontifex et al., 2019).

To characterize the sensitivity of this secondary analysis, we conducted post-hoc simulation-based power analysis, which appropriately accounts for the nested crossover structure and crossed random effects (Johnson et al., 2015). Specifically, we generated 1,000 new datasets by drawing from the estimated parameter distributions of the fitted mixed-effects model and refitting the full model to each simulated dataset with SIMR r package approach (Green & MacLeod, 2016). We tested both the strongest and weakest significant C-HIIT × brain volume interactions across model families, providing a conservative bound. Power exceeded 99% across all four models at n = 13, with 100% convergence in all simulations. Power curve analysis (subsampling at n = 5, 7, 9, 11, 13) confirmed that power remained ≥80% at n ≥ 9 in all cases, indicating that the study was adequately powered to detect the interaction effects on which the conclusions rest, even under the most conservative scenario.

### 2.2 Experimental conditions and procedures of within-subject cross-over trial

Participants completed three randomized experimental conditions with two-week washout periods under supervision at the IRyS laboratory. In the Sedentary Condition (SC), adolescents watched a nature documentary for 90 minutes indoors (20-22°C, standardized lighting). In the Moderate-Intensity Continuous Training (MICT) condition, participants engaged in outdoor self-paced brisk walking targeting ~60% of estimated maximum heart rate (Butte et al., 2018). In the Cooperative High-Intensity Interval Training (C-HIIT) condition, participants performed four sets of four cooperative exercises (20:40-second work-to-rest ratio) targeting ≥85% of estimated HRmax (Chen et al., 2014).

All physical activity sessions included 4-minute warm-up and cool-down periods. Heart rate was monitored using Polar H10 monitors, with maximum heart rate calculated as 220 minus age. Energy expenditure was matched across conditions using BMR estimates and MET values (SC: 1.3, MICT: 4.9, C-HIIT: 7.4) to ensure comparable physiological demands through different modalities (Butte et al., 2018).

Reading evaluations occurred ~20 minutes post-intervention. Session order was randomized, with sessions held every two weeks on the same weekday after school. The study design is presented in the CONSORT flow chart in Figure 1. A typical measurement and intervention session is described in the Supplementary Material.

**Figure 1.**
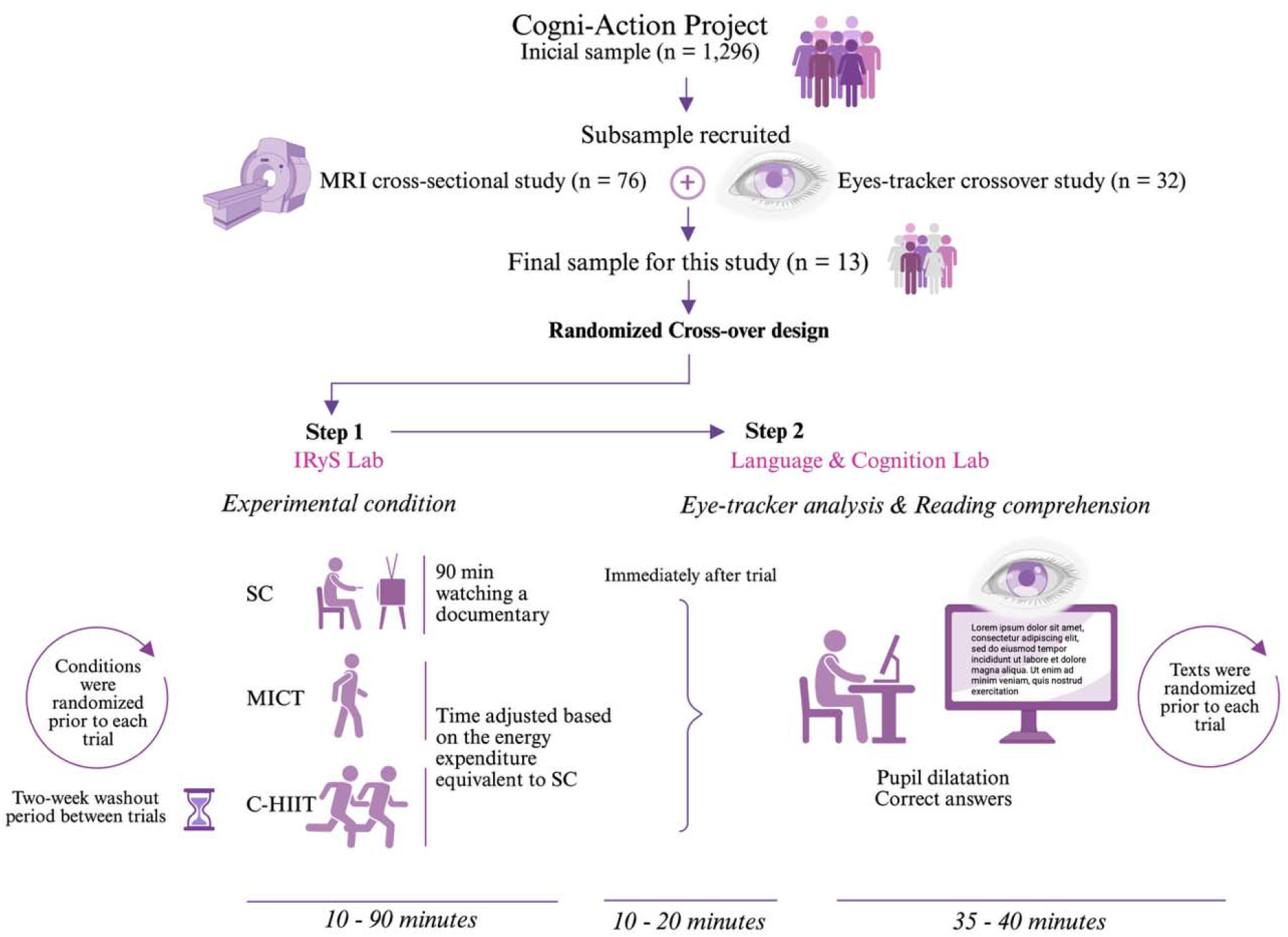
CONSORT-based flowchart of study design and procedures. SC: Sedentary Condition. MICT: Moderate-Intensity Continuous Training. C-HIIT: Cooperative High-Intensity Interval Training.

**Figure 2.**
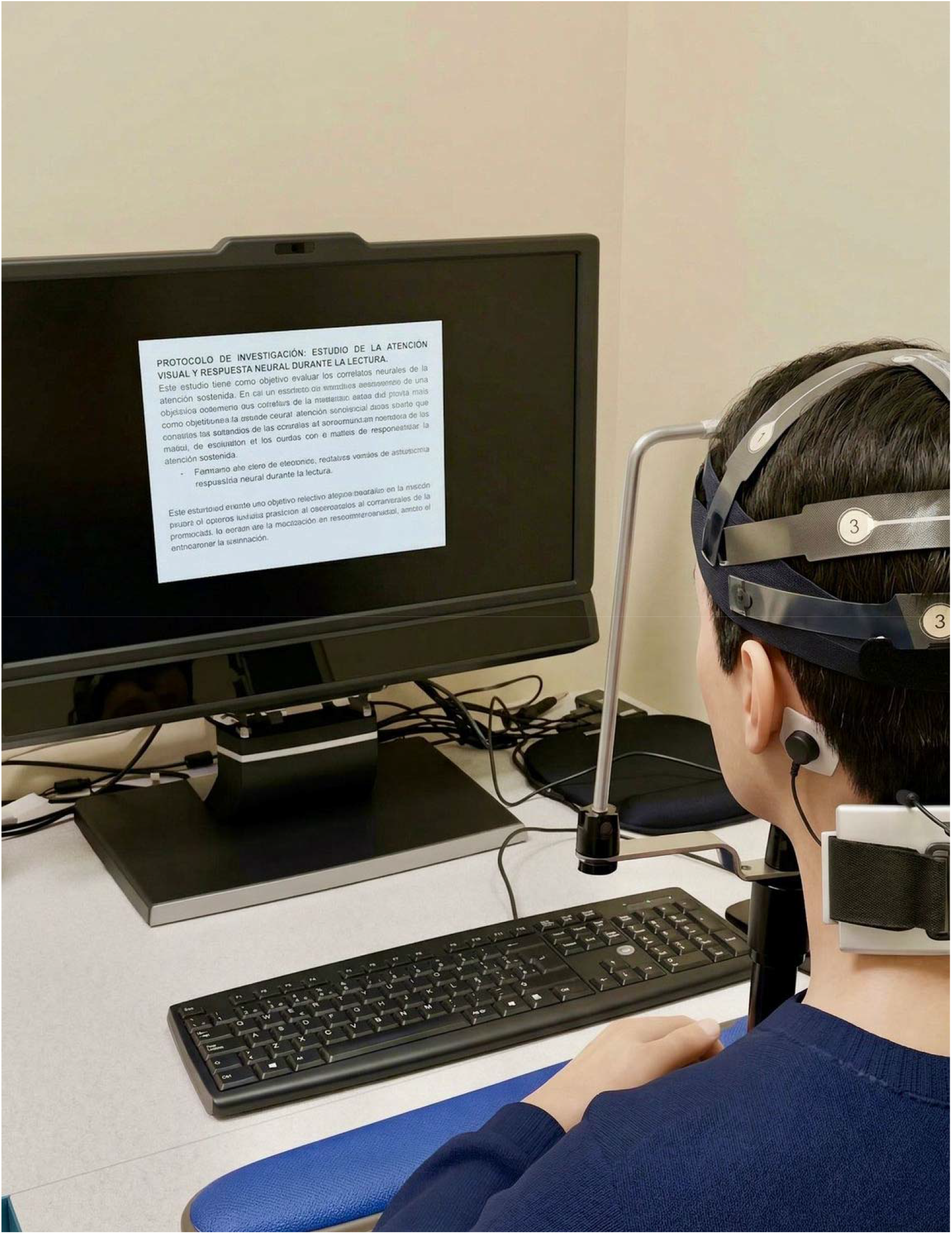
Setup of the reading task with head stabilization for data recording. To ensure full subject anonymity and comply with strict ethical guidelines, the visible human figure has been computer-generated instead of using insufficient methods such as pixelation.

### 2.3 Cognitive Load and Comprehension Measurements

Participants read three randomized narrative texts in Spanish (Greek and African myths) with similar syntactic complexity and word counts (506-555 words). Pupil dilation during reading served as a physiological proxy for CL (Engelhardt et al., 2010; Zou et al., 2023), captured using a Tobii TX-300 eye tracker (300 Hz sampling rate, monocular tracking). All assessments occurred under identical lighting conditions (Lux: 698.67±106.9). Participants were medication-free, instructed to abstain from caffeine for 24 hours, and obtain ≥8 hours of sleep prior to testing.

Comprehension was assessed with seven multiple-choice questions per text, validated by three specialists (Fleiss’ Kappa=0.83). Internal consistency was acceptable (Cronbach’s α=0.76) based on pilot testing.

### 2.4 MRI acquisition and processing

T1-weighted magnetization-prepared rapid gradient echo (MPRAGE) images were processed using FreeSurfer 7.4.1 on Neurodesk via the cross-sectional recon-all pipeline (e.g., recon-all -s subject_id -i T1image -all). From the outputs we extracted gray-matter volumes for the left and right hippocampi, the left pars opercularis, and the brainstem. Hippocampal and brainstem volumes were obtained from the aseg.stats file, whereas the left pars opercularis was extracted from the lh.aparc.stats file; both aseg and aparc outputs are based on the Desikan–Killiany atlas. These regions were selected based on three convergent criteria: (1) established roles as strong neuroanatomical predictors of reading comprehension performance (Keller et al., 2024; Skoe et al., 2017), (2) documented sensitivity to acute physical activity through noradrenergic pathways that enhance cognitive function (Plini et al., 2023; Tsetsenis et al., 2022; Yebra et al., 2019), and (3) functional specialization across different levels of the construction-integration reading model, with the pars opercularis supporting phonological-orthographic and semantic processing (surface code/textbase), the hippocampus enabling contextual integration and situation model construction, and the brainstem (particularly locus coeruleus) regulating arousal, attention, and pupillary responses that index cognitive load (Basinger & Hogg, 2025; Keller et al., 2024; Kintsch, 1988). Quality control was independently performed following the ENIGMA Consortium Cortical Quality Control (QC) Protocol 2.0 (https://enigma.ini.usc.edu), which included visual inspection of segmentation accuracy, detection of outliers using statistical thresholds (>2.5 standard deviations from the mean), assessment of motion artifacts, and verification of anatomical plausibility of extracted volumes. This processing pipeline provided precise neuroanatomical metrics essential for assessing the structural correlates of reading comprehension in our study.

### 2.5 Statical analysis

First, due to the large differences in scale among pupil dilation, reading comprehension scores, and gray matter brain volumes (expressed in cubic millimeters), all continuous variables were standardized using Z-scores (mean-centered and divided by the standard deviation) to facilitate interpretability and comparability across measures. Data normality was evaluated using Shapiro–Wilk tests, inspection of residual distributions, and Q–Q plots.

To account for the repeated-measures cross-over design and the hierarchical structure of the data, analyses were conducted using mixed-effects modeling. Pupil dilation (CL) was analyzed using linear mixed-effects models, whereas reading comprehension performance (number of correct answers) was analyzed using generalized linear mixed-effects models with a Poisson distribution and log link function. All models were implemented using the lme4 package in R (Version 4.0.1; R Core Team, 2021).

Experimental condition (PA) was specified as a fixed effect. Participants and texts were included as random intercepts to account for inter-individual variability and text-specific differences in reading difficulty, thereby avoiding the language-as-fixed-effect fallacy. In line with current recommendations for confirmatory mixed-effects modeling, a maximal random-effects structure justified by the experimental design was adopted to control Type I error at the model level and to appropriately model dependency among repeated observations (Barr et al., 2013).

Effect modulation by gray matter brain volume was examined by including a condition × brain volume interaction term in the mixed-effects models. Brain volume was treated as a continuous moderator to preserve variability and maximize statistical power. A significant interaction indicates that the effect of physical activity condition on pupil dilation or reading comprehension varies as a function of baseline gray matter volume.

For generalized mixed-effects models, results are reported as incidence rate ratios (IRRs), with IRRs greater than 1 indicating proportional increases and IRRs less than 1 indicating proportional decreases relative to the reference condition. Linear mixed-effects model results are reported as regression coefficients (β). Given the modest sample size (n = 13), statistical significance was evaluated using permutation-based p-values rather than asymptotic approximations. Condition labels were permuted within each participant across 10,000 iterations, preserving the crossover structure of the data, and two-tailed p-values were computed as the proportion of permuted statistics meeting or exceeding the absolute value of the observed statistic, with a minimum attainable p-value of 0.0001. Fixed effects estimates and standard errors are reported from the mixed-effects models, as these remain the optimal estimators given the hierarchical design (Corraini et al., 2017)

No additional covariates were included for three complementary reasons. First, the within-subjects crossover design inherently controls for all stable between-participant characteristics (including baseline fitness, cognitive capacity, and anthropometric variables) as each participant serves as their own control across conditions; these sources of variance are absorbed by the participant random intercept. Second, condition-level variables that could vary between sessions were experimentally standardized: the interval between exercise and cognitive testing was fixed at approximately 20 minutes for all participants and conditions, session order was randomized, texts were counterbalanced, and testing environment was held constant. Third, model parsimony was prioritized given the sample size: including additional covariates with n = 13 would introduce overfitting and convergence instability without improving causal inference, given that the design already eliminates the confounds they would target.

## 3. Results

Descriptive analyses characterized the sample’s baseline values (Table 1). No participants were at risk of excessive abdominal fat or obesity. The waist-to-height ratio was used as a key indicator due to its stronger agreement with DEXA-measured fat mass (the gold standard) compared to BMI and its relevance to obesity’s negative impact on cognitive performance. BMI also was computed for comparison in SD according to the age of our sample. The volume values of the areas of interest, expressed in cubic millimeters, are presented in Table 1.

**Table 1.** Participants characteristics.

| Indicator | Participants Mean ± SD |
| --- | --- |
| Age (years) | 12.0 ± 0.7 |
| Body Weight (kg) | 54.6 ± 12.1 |
| Body Height (cm) | 156 ± 10.3 |
| BMI (SD) | 1.37 ± 1.01 |
| Waist-to-height ratio | 0.45 ± 0.05 |
| Correct Answers SC | 3.85 ± 0.81 |
| Correct Answers MICT | 4.37 ± 1.39 |
| Correct Answers C-HIIT | 4.49 ± 1.09 |
| Left hippocampus volume (mm<sup>3</sup>) | 3854 ± 518 |
| Right hippocampus volume (mm<sup>3</sup>) | 4074 ± 398 |
| Left pars opercularis volume (mm<sup>3</sup>) | 5852 ± 767 |
| Brainstem (mm<sup>3</sup>) | 20517 ± 1106 |

kg: kilograms. Cm: centimeters. SD: standard deviation. mm<sup>3</sup>: cubic millimeters

Mixed-effects models fitted for pupil diameter demonstrated conditional R^2^ values between 0.80-0.85 across all analyses, indicating that the combination of fixed and random effects explained a substantial proportion of total variability. The high ICC values (ranging from 0.80-0.82 across all models) confirm that systematic differences between participants and items were effectively captured by the random structure, reflecting the within-subjects crossover design where participants served as their own controls (Baayen et al., 2008; Pontifex et al., 2019). In all models, physical activity was a significant predictor of CL and reading comprehension. However, the baseline brain volume was not significant in any model of CL and only in two models of reading comprehension, in pars opercularis (IRR=0.83, 95% CI [0.79, 0.87] p<0.001) and brainstem (IRR=0.90, 95% CI [0.85, 0.95], p<0.001).

### Cognitive Load

Mixed-effects models revealed significant differences in CL between physical activity conditions across all examined brain regions. Without considering brain volume modulation, principal effects show MICT consistently demonstrated higher CL (pupil dilatation) than the SC across all models, while C-HIIT showed lower CL (pupil contraction) compared to MICT. The pattern for C-HIIT versus SC varied by brain region, with some regions showing significant differences only in models of pars opercularis and brainstem.

When brain volume was incorporated as a baseline modulator as interaction term, the relationships between conditions changed substantially. The volume interactions revealed enhanced CL responses (pupil dilatation) for C-HIIT compared to SC in hippocampal regions, while reducing the CL (pupil contraction) responses for both physical activities conditions in pars opercularis and brainstem regions. These modulation effects maintained the relative pattern of C-HIIT showing lower CL (pupil contraction) than MICT across all regions.

### Reading Comprehension

Both physical activity conditions consistently outperformed the SC across all brain regions. Principal effects showed than C-HIIT was superior performance compared to MICT in hippocampal regions and brainstem, while performance was equivalent between exercise conditions in pars opercularis.

Brain volume modulation as interaction term fundamentally altered these relationships. Volume interactions reduced MICT benefits while enhancing C-HIIT advantages, creating substantial separation between physical activity conditions. In hippocampal regions, volume modulation eliminated MICT superiority over sedentary controls while amplifying C-HIIT benefits. In pars opercularis, where physical activity conditions were initially equivalent, volume modulation specifically enhanced C-HIIT performance, creating a significant advantage. The brainstem showed the strongest modulation effect, with C-HIIT demonstrating markedly superior performance over both other conditions.

Figures 3 and 4 show all models without and with interaction to compare the modulation effects of brain volume. Complete statistical details including confidence intervals and p-values are provided in Supplementary Material.

**Figure 3.**
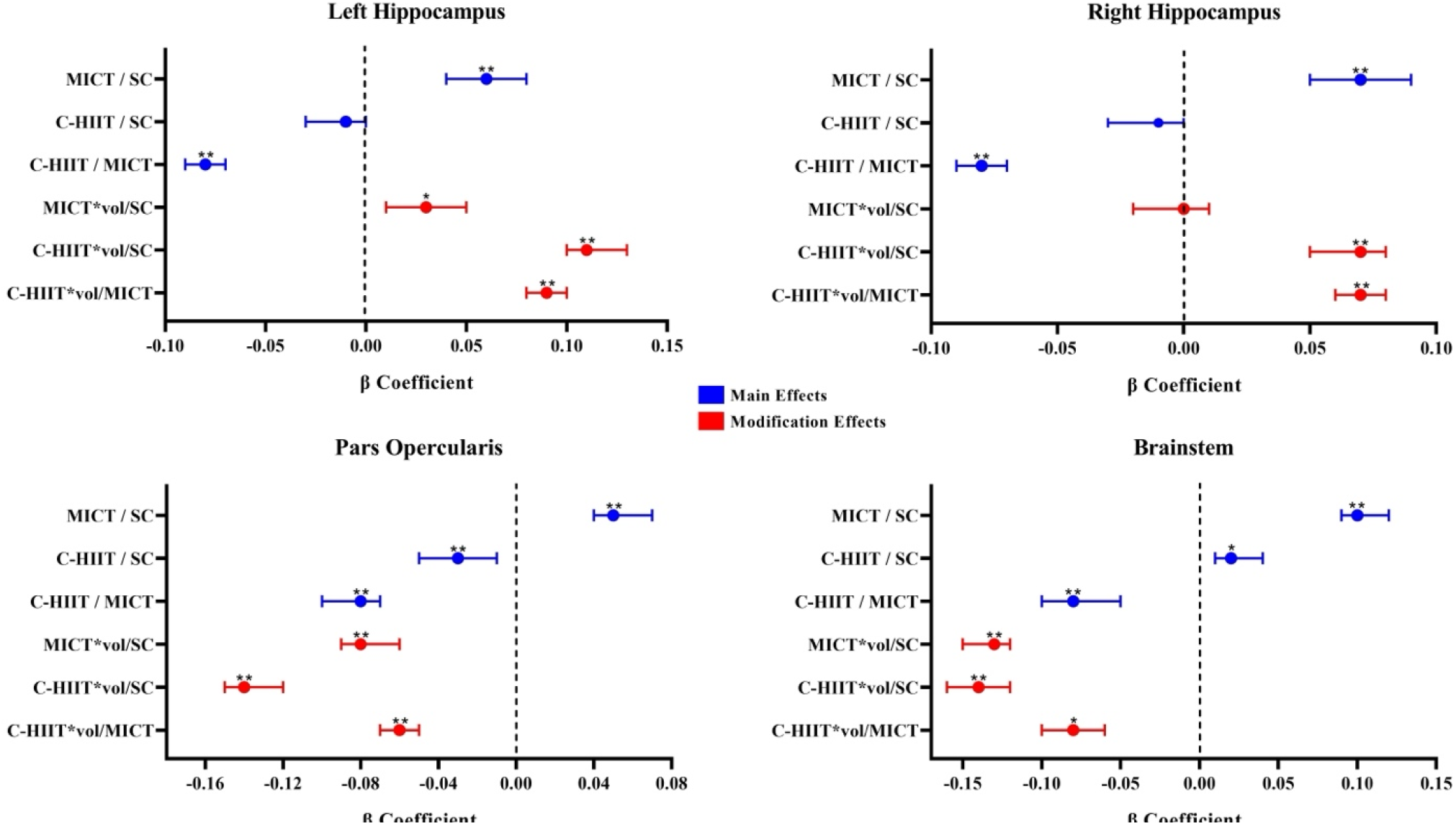
Comparison of brain volume on modulating cognitive load. Data are presented as comparator vs reference. MICT: Moderate-Intensity Continuous Training. SC: Sedentary Condition. C-HIIT: Cooperative High-Intensity Interval Training. **:<0.001. *:<0.05

**Figure 4.**
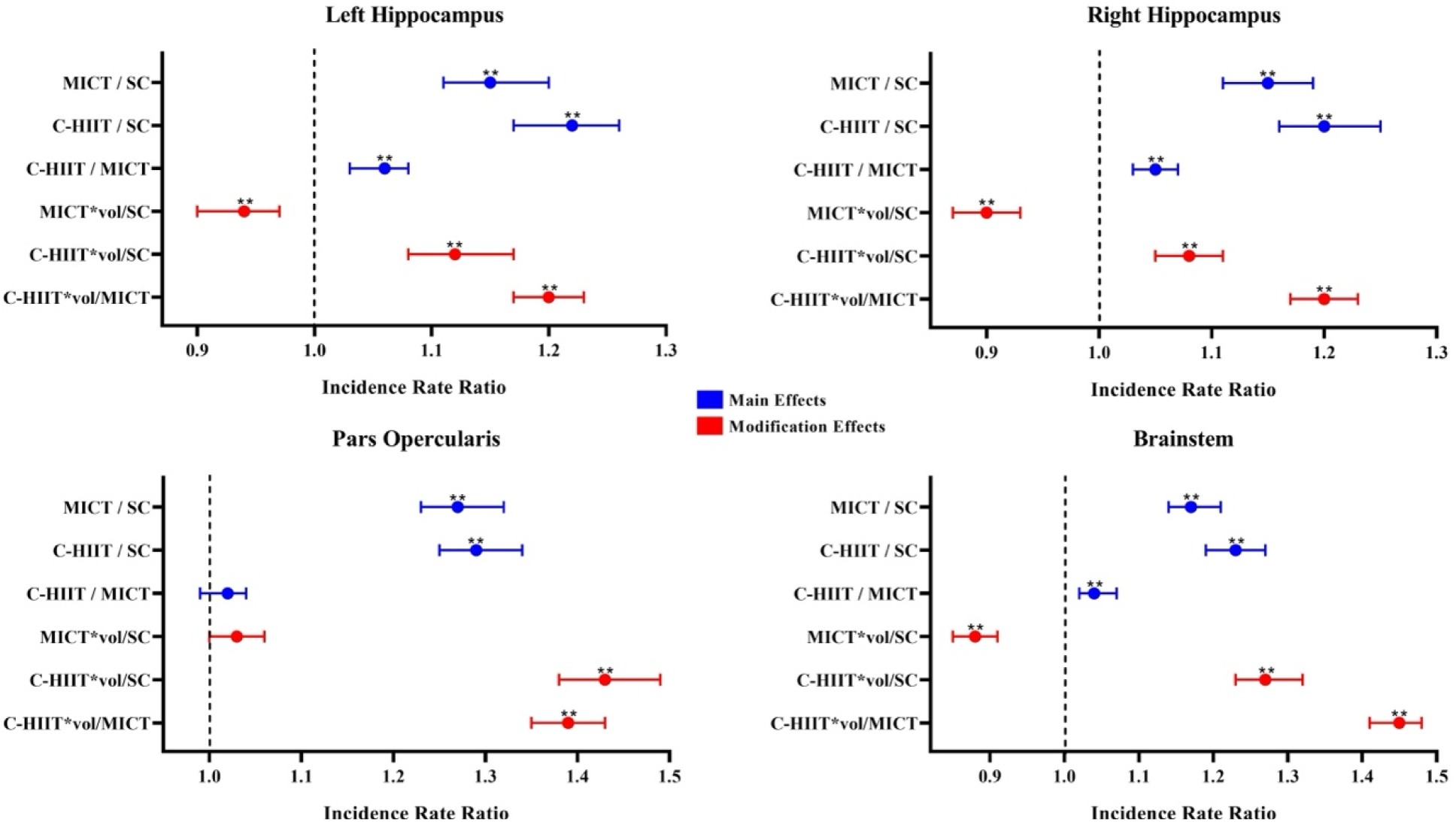
Comparison of brain volume on modulating reading comprehension. Data are presented as comparator vs reference. MICT: Moderate-Intensity Continuous Training. SC: Sedentary Condition. C-HIIT: Cooperative High-Intensity Interval Training. **:<0.001. *:<0.05

## 4. Discussion

This study aimed to determine how brain volume of specific regions could modulate the effect of physical activity on reading comprehension and the associated CL, indexed by pupil dilation. A key finding across our models is that gray volume in each of the assessed brain structures was not a significant predictor in any model of CL and only in two models of reading comprehension. This suggests that pre-existing neuroanatomy in regions associated with reading comprehension and visual attention does not, on its own, significantly predict CL and only partially predict reading comprehension in schoolers. Instead, it was the interaction between gray matter volume and physical activity that altered the influence of active conditions on reading comprehension and associated CL. Notably, physical activity alone remained a significant predictor in all models, even without accounting for modulation effects. Thereby, depending on the specific brain structure examined, our analyses revealed mixed results, highlighting the importance of consider individual neuroanatomical differences. However, a consistent observation was that school students participating in C-HIIT showed improved reading comprehension and different pattern of CL according to brain structure, compare with SC and MICT.

### 4.1 Physical activity effect on Reading comprehension and CL

The type of physical activity was fundamental in the findings encountered. When brain volume was considered only C-HIIT demonstrated superiority SC but also MICT, registering a greater number of correct responses with a different and apparently superior distribution of cognitive resources. This superiority was demonstrated through variations in pupil dilation that indicated differential CL levels in real time during reading, while simultaneously maintaining superior performance on comprehension questions. These findings reflect distinct patterns of reading processing, or potentially even different levels of cognitive processing depth, depending on the specific brain structures examined.

During reading, different brain processes are generated as explained by the construction-integration model (Kintsch, 1988). This model postulates that comprehension involves: (1) surface code representation (literal form of the text), (2) textbase (propositions and explicit relationships), and (3) construction of the situation model (contextual and relational representation of content), through phases of massive construction of activations followed by selective integration that stabilizes the final representation, progressing from less to more complex process (Kintsch, 1988). In this sense, a possible explanation of our results is grounded in the brain structures involved in this process, where the pars opercularis and brainstem may be involved in surface code and textbase processes, while the hippocampus intervenes in the situation model since all these brain structures are strong predictor of reading comprehension (Keller et al., 2024).

On one hand, the pars opercularis is a key node in phonological-orthographic processing and in subvocal recoding, functions directly involved in lexical recognition and initial propositional assembly of the text (Glezer et al., 2016) with a critical role in language and semantic processing (Keller et al., 2024). Likewise, the brainstem, particularly the locus coeruleus, is key in top-down attentional processes that support initial phases of information processing (Katsuki & Constantinidis, 2014). On the other hand, the hippocampus is related to information integration processes and mental constructions based on prior knowledge and contextual information (Hausser et al., 2021; Yebra et al., 2019). In this sense, acute physical activity was associated with differential response of these structures, particularly C-HIIT according to our results, and these effects appear linked to enhanced noradrenergic signaling mechanisms. A possible neurochemical explanation, consistent with animal and human evidence, is that during physical activity, the excitability of the locus coeruleus (houses in brainstem) increases, which projects massively through different neural connections throughout the brain and generates an increase in brain catecholamines (McMorris, 2021), one of its main neurotransmitters, which projects to the prefrontal cortex, the inferior frontal gyrus (where the pars opercularis is located), and the hippocampus, generating greater efficiency in processing speed, information integration and inhibition, improving top-down attentional control (Hansen et al., 2024; Tsetsenis et al., 2022). Likewise, possibly the lexical processing dependent on the pars opercularis is also favored by this increase in catecholamines (Hansen et al., 2024). Furthermore, this noradrenergic response improves associative encoding in the hippocampus, optimizing complex information integration processes (Yebra et al., 2019). In this sense, it has been observed that noradrenaline discharges (which can be produced by physical activity) can increase or decrease neuronal excitability in response to stimuli, improving coordination of different brain and synaptic areas, modulating theta and gamma oscillations to selectively synchronize activity between brain regions and improving information processing (Guedj et al., 2017; Tully et al., 2007). However, these mechanisms were not directly measured in the present study and remain as an speculative hypothesis pending neurochemical or neuroimaging confirmation.

Based on these mechanisms, our results indicate that C-HIIT was associated with lower pupil dilation (reflecting reduced CL) during reading while maintaining a higher rate of correct responses, suggesting enhanced comprehension with lower “cognitive cost”. When modulation by the brainstem and pars opercularis is considered, this pattern is consistent with more efficient allocation of cognitive resources in less complex stages of reading (surface code and textbase). The pars opercularis, in particular, plays a critical role: previous literature identifies it as one of the strongest predictors of reading comprehension due to its central involvement in semantic and language processing (Keller et al., 2024), which are essential components of surface code and textbase within the integration– construction model (Kintsch, 1988). In this sense, enhanced sustained attention associated with brainstem activation (Hansen et al., 2024), together with more efficient semantic processing, may be associated with better coupling between surface code and textbase with hippocampal activity and the situation model. This dynamic allows resources to be preserved for more complex processes of integration and encoding in the hippocampus, which demand higher CL (Keller et al., 2024). The varied patterns across regions reflect functional specialization: brainstem and pars opercularis (foundational processing) showed volume-dependent CL reductions under C-HIIT, while hippocampus (higher-order integration) demonstrated volume-dependent comprehension enhancements (Keller et al., 2024; Kintsch, 1988). Critically, baseline volume predicted outcomes only when interacting with physical activity, suggesting neuroanatomical capacity requires appropriate physiological activation to manifest functionally. C-HIIT’s superior outcomes are consistent with greater locus coeruleus activation and phasic noradrenergic bursts, as suggested by prior evidence (Vazey et al., 2018), whereas MICT’s sustained moderate intensity may be insufficient to fully engage these neurochemical pathways to the same degree, though this cannot be determined from the present data.

Regarding the specific difference between MICT vs C-HIIT, while previous evidence has indicated that both types of physical activity have important benefits for cognition (Festa et al., 2023), the cyclical nature of C-HIIT may generate an improvement compared to MICT. C-HIIT may generate repeated perturbations in homeostasis that may enhance lactate shuttling to the brain and, in this sense, physical activity-enhanced brain executive function is related to cerebral lactate metabolism (Xue et al., 2022). Likewise, the greater intensity of C-HIIT may be associated with a greater brain noradrenergic response compared to MICT. Animal models have shown that higher intensity PA generates greater brain catecholamine release, increasing cognitive function (Arbat-Plana et al., 2019; Basso & Suzuki, 2016). This noradrenergic response has been proposed to modulate other neurotransmitters such as glutamate, dopamine, and serotonin, key to hippocampal function (Moreno-Castilla et al., 2017; Song & Tan, 2023), and possibly improving the complete construction-integration process.

### 4.2 Strengths and limitations

This study has several methodological strengths. The crossover design, which is the most rigorous one when investing the effects of acute PA on cognition (Pontifex et al., 2019), minimized interindividual variability in responses to PA and the cognitive demands of the texts. To our best knowledge, our project was the first to examine PA’s effect on reading comprehension and CL. In the same way this study is also the first to assess how gray matter volume in principal reading predictor regions regions—bilateral hippocampus, and left pars opercularis and brainstem—modulates PA’s effect on CL during reading and reading comprehension. Finally, this study contributes valuable data on an underrepresented population (Latin American students) in scientific research.

However, important limitations must be acknowledged. The modest sample size (n=13) limits power and generalizability, requiring replication in larger samples. The male-only sample prevents examining sex differences in brain volume × exercise interactions. We cannot disentangle intensity from modality effects as C-HIIT differs from MICT in both dimensions. Age-predicted HRmax formulas lack individual precision. Cross-sectional brain volume measurement precludes causal inferences about structural differences. The indoor-outdoor environmental contrast introduces potential confounds beyond exercise effects.

## 5. Conclusion

This study provides novel evidence that gray matter volume in brain regions critical for reading comprehension does not independently predict CL or reading comprehension in school adolescents. Instead, these neuroanatomical individual differences interact with physical activity modalities to shape learning outcomes, highlighting the need for personalized educational strategies that account for brain variability in school settings. The synergistic interaction between individual neuroanatomy and physical activity modality emerges as the determining factor for cognitive outcomes. C-HIIT demonstrated superior effects on both CL and reading comprehension compared to MICT and SC, with brain volume interactions consistently enhancing C-HIIT benefits while attenuating MICT effects across all examined regions.

The superior performance of C-HIIT is consistent with more efficient cognitive processing during reading, as indexed by lower pupil dilation alongside higher comprehension accuracy. These findings demonstrate that physical activity modality selection is critical for maximizing cognitive benefits in adolescents. These findings highlight the potential relevance of considering both physical activity modality and individual brain morphology when examining cognitive outcomes in school settings.

## Supporting information

Supplementary results

## Data Availability Statement

The data presented in this study are available on request from the corresponding author. The data are not publicly available due to ethical concerns.

## Funding

This “Cogni-Action Project” was supported by the National Commission for Scientific and Technological Research CONICYT/FONDECYT INICIACION 2016 grant no. 11160703.

## Conflicts of Interest

The authors declare no conflicts of interest.

## Acknowledgement

Ricardo Martínez-Flores is supported by the National Agency for Research and Development (ANID)/Scholarship Program/DOCTORADO BECAS CHILE/2024–(Grant Nº 72240103).

We wish to thank the entire team that made this work possible, starting with the students and parents, the administration of Ruben Castro School, as well as the technical staff and researchers, including Felipe Porras, Inti Federici, Constantino Dragicevic, Aland Astudillo, and Eduardo Méndez. We also want to acknowledge the Independent Imagenology Center Quintaimagen, Viña del Mar, Chile. We extend special recognition to the late Dr. Giovani Parodi, who opened the doors of his laboratory and trusted us to carry out this work.

## Declaration of generative AI and AI-assisted technologies in the manuscript preparation process

During the preparation of this work the author(s) used ChatGPT-4o in order to improve language and readability. After using this tool/service, the author(s) reviewed and edited the content as needed and take(s) full responsibility for the content of the publication.

## Author contributions

Both R.M-F and C.C-M contributed equally. The corresponding author (C.C-M) attests that all listed authors meet authorship criteria. R.M-F, and C.C-M they carried out the conceptualization of the study. R.M-F and conducted statistical analysis. R.M-F and C.C-M wrote the original draft. H.S, J.S-M, P.S-U, B.T, F.H, F.P, M.M and L.Z, performed a critical revision and editing of the manuscript. All authors have read and agreed to the published version of the manuscript.

## Ethical approval statement

All the procedures performed in this study involving human participants were in accordance with the ethical standards of the institutional and/or national research committee and with the 1964 Helsinki declaration and its later amendments or comparable ethical standards. The study was approved by the Bioethics Committee of the Pontificia Universidad Católica de Valparaíso (BIOEPUCV-H103–2016).

## Patient consent statement

Written informed consent was obtained from the school principal and parents, along with the participants’ assent.

## Notes

### Competing Interest Statement

The authors have declared no competing interest.

