## Supplementary results for "Gray matter Volume Modulates the Effect of Acute Physical Activity on Reading Comprehension and Cognitive Load in Adolescents –The Cogni-Action Project"

*S1. Experimental conditions and procedures description of within-subject cross-over trial*

Participants completed three experimental conditions with two weeks of wash out periods between trials in a randomized order at the IRyS Laboratory, always under the supervision of a trained research assistant. In the Sedentary Condition (SC), adolescents watched a nature documentary for 90 minutes in a controlled indoor environment emulating at certain point a regular school class (20–22 °C, standardized lighting, and absence of environmental noise). To ensure participants remained attentive, the assistant periodically asked simple content-related questions.

In the Moderate-Intensity Continuous Training (MICT) condition, participants engaged in a self-paced brisk walk outdoors, targeting approximately 60% of their estimated maximum heart rate (Butte et al., 2018). In the Cooperative High-Intensity Interval Training (C-HIIT) condition, participants performed four sets of four cooperative exercises designed to engage cardiorespiratory, speed-agility, and coordinative demands (Chen et al., 2014). These activities followed a 20:40-second work-to-rest ratio and aimed to reach at least 85% of estimated HRmax. The term cooperative exercises refers to cognitively and physically demanding tasks that require interpersonal interaction, coordination, and rapid decision-making (Chen et al., 2014).

All physical activity sessions included a 4-minute warm-up (running, lateral movements, dynamic stretching) and a 4-minute cool-down. Physical activity intensity was monitored using Polar H10 heart rate monitors. Maximum heart rate was calculated using the 220 minus age formula and effort calculated as (exercise HR/[220–age])×100. Although heart rate–based formulas are widely used in children and adolescents, they may lack individual precision; we applied such a formula to ensure comparability with the large body of prior studies in similar populations (Cicone et al., 2019). Nevertheless, this constitutes a limitation of our study, and future research should consider more accurate approaches for estimating exercise intensity (e.g., graded exercise testing) (Cicone et al., 2019).

Both physical activity conditions (MICT and C-HIIT) were performed outdoors to emulate typical school recess environments, whereas the SC condition was carried out indoors to mirror a classroom setting. This indoor–outdoor contrast captures ecologically valid differences in adolescents’ daily activity contexts and, rather than representing a methodological weakness, should be regarded as an important contextual factor to acknowledge when interpreting cognitive outcomes (Herold et al., 2019; Mavilidi et al., 2025).

Energy expenditure was matched across all three conditions to ensure comparable physiological demands delivered through different modalities. Specifically, basal metabolic rate (BMR) was estimated for each participant, and energy expenditure during the SC was calculated as METS (1.3) × BMR (kcal/min) × duration (90 minutes). For MICT and C-HIIT, the duration of physical activity was tailored to match this estimated expenditure using their respective MET values (4.9 and 7.4, respectively) (Butte et al., 2018). For detailed methodology on energy matching procedures, please refer to a previous published study (Martínez-Flores et al., 2025).

Each participant attended three sessions, scheduled every two weeks to allow a wash-out period, with session order randomized and held on the same weekday after school. A typical session involved: (1) Arrival at the IRyS Laboratory for anthropometric measurements and randomization into the physical activity protocol (duration: 10–90 minutes); (2) execution of the assigned physical activity intervention; (3) escorted to the "Language & Cognition" Laboratory for a reading evaluation initiated within 20 minutes of physical activity completion (lasting 35–40 minutes); and (4) a comprehension test of seven questions (3–5 minutes) before being escorted back to school. The reading texts were randomized across sessions, such that all participants completed the three physical activity protocols and read the three texts under different conditions. For more information on the experimental protocol, please refer to a previous published study (Martínez-Flores et al., 2025).

*S2. Cognitive Load and Comprehension Measurements*

In the Lenguaje & Cognición Laboratory at the authors' institution, participants read three randomized narrative texts in Spanish (Greek and African myths) designed by linguistic experts. All texts had similar syntactic complexity and word counts (506–555 words) to minimize learning effects.

Previous studies have established that pupil dilation is a reliable physiological indicator of CL during reading, particularly in response to syntactic complexity, lexical difficulty, and discourse integration demands (Engelhardt et al., 2010; Zou et al., 2023). In our study, we did not include a separate behavioral task to assess CL; instead, we used pupil dilation during reading as an established proxy for moment-to-moment CL (van der Wel & van Steenbergen, 2018; Zou et al., 2023).

To ensure consistency in pupil measurement, all participants were assessed under identical conditions and lighting environments (Lux mean ± SD of 3 measures 698.67± 106.9). Furthermore, participants were medication-free and were instructed to abstain from caffeine consumption for 24 hours before testing and to obtain a minimum of 8 hours of sleep the night prior to the experimental sessions, according to guidelines for pupilography (Kelbsch et al., 2019). The texts were carefully designed to correspond with students' curricular knowledge and the subjects they were studying at their educational level, minimizing the potential confounding effects of arousal or luminance-induced pupil changes. Pupil dilatation as proxy of CL was captured using the Tobii TX-300 eye tracker, operating at a sampling rate of 300 Hz with monocular tracking. Stimuli were presented on a screen with a resolution of 1920 × 1080 pixels. The experiment was constructed, data collected, and areas of interest (i.e., individual words) defined using Tobii Studio Pro software. Reading evaluations began about 20 minutes after each experimental condition, simulating a typical school recess, and comprehension was assessed with seven questions per text. The test consisted of multiple-choice questions with a single correct option, enabling an objective and standardized evaluation process. To validate the content, three specialists in reading instruction and educational assessment were consulted. They examined each item to ensure clarity, relevance, and alignment with the targeted comprehension level. The level of inter-rater agreement was assessed using the Fleiss' Kappa index, yielding a value of 0.83, which reflects a high degree of concordance among the experts. Due to the exploratory nature of the reading analysis, the number of questions was intentionally limited.

To assess the reliability of the reading comprehension questions, a pilot study was conducted with a group in a subsample of students from the original crossover trial sample who shared similar characteristics with the participants included in the final analysis. The responses were analyzed using Cronbach's alpha coefficient, resulting in a value of 0.76. This indicates an acceptable level of internal consistency for a test of this nature, particularly given the limited size of the pilot sample. Text validation procedures, including syntactic complexity criteria and readability assessments, were conducted as part of the material development process. For more detailed information on the eye tracking measurement, including processing of the data, and the text used, please refer to a previous published study (Martínez-Flores et al., 2025).

*S3.* *Cognitive Load* *Results - Full Description*
Mixed effect models revealed significant differences in CL between all three conditions across all brain regions examined. In the left hippocampus model, MICT showed greater CL than SC (β=0.06, p<0.001, 95% CI [0.04, 0.08]), while C-HIIT vs SC was not significant. C-HIIT showed significantly lower CL than MICT (β=-0.08, p<0.001, 95% CI [-0.09, -0.07]). When left hippocampus volume was considered as a baseline modulator, significant interactions emerged for MICT vs SC (β=0.03, p=0.003, 95% CI [0.01, 0.05]), C-HIIT vs SC (β=0.11, p<0.001, 95% CI [0.10, 0.13]), and C-HIIT vs MICT (β=0.09, p<0.001, 95% CI [0.08, 0.10]). This indicates that both physical activity conditions showed greater CL than SC when volume interactions were considered, and the MICT vs C-HIIT relationship was reversed, generating greater CL in C-HIIT.

In the right hippocampus model, MICT demonstrated greater CL than SC (β=0.07, p<0.001, 95% CI [0.05, 0.09]), while C-HIIT vs SC was not significant. C-HIIT showed significantly lower CL than MICT (β=-0.08, p<0.001, 95% CI [-0.09, -0.07]). When right hippocampus volume was considered as a baseline modulator, the MICT vs SC interaction was not significant, while both C-HIIT vs SC (β=0.07, p<0.001, 95% CI [0.05, 0.08]) and C-HIIT vs MICT (β=0.07, p<0.001, 95% CI [0.06, 0.08]) showed significant interactions. This suggests that volume modulation primarily affected C-HIIT responses, making C-HIIT show greater CL than SC and reducing the difference with MICT.

In the pars opercularis model, MICT showed increased CL compared to SC (β=0.05, p<0.001, 95% CI [0.04, 0.07]), while C-HIIT showed reduced CL compared to SC (β=-0.03, p<0.001, 95% CI [-0.05, -0.01]). C-HIIT demonstrated significantly lower CL than MICT (β=-0.08, p<0.001, 95% CI [-0.10, -0.07]). When pars opercularis volume was considered as a baseline modulator, significant interactions emerged for MICT vs SC (β=-0.08, p<0.001, 95% CI [-0.09, -0.06]), C-HIIT vs SC (β=-0.14, p<0.001, 95% CI [-0.15, -0.12]), and C-HIIT vs MICT (β=-0.06, p<0.001, 95% CI [-0.07, -0.05]). This indicates that volume modulation attenuated the CL responses for both physical activity conditions while maintaining the relative pattern of MICT > C-HIIT in terms of CL.

In the brainstem model, MICT showed the highest CL compared to SC (β=0.10, p<0.001, 95% CI [0.09, 0.12]), while C-HIIT also showed increased CL compared to SC (β=0.02, p=0.012, 95% CI [0.01, 0.04]). C-HIIT showed significantly lower CL than MICT (β=-0.08, p<0.001, 95% CI [-0.10, -0.05]). When brainstem volume was considered as a baseline modulator, MICT vs SC showed significant interaction (β=-0.13, p<0.001, 95% CI [-0.15, -0.12]), C-HIIT vs SC showed significant interaction (β=-0.14, p<0.001, 95% CI [-0.16, -0.12]), and C-HIIT vs MICT showed significant interaction (β =-0.08, p=0.025, 95% CI [-0.10, -0.06]). This suggests that volume modulation substantially reduced CL responses for both physical activity conditions while maintaining the relative pattern of reduce C-HIIT condition in terms of CL compared to MICT, also observed in pars opercularis.

*S3.* *Reading Comprehension Results - Full Description*

For reading comprehension, both PA conditions consistently showed higher response rates compared to SC across all brain regions, however when brain volume was considered only C-HIIT was superior to SC and also to MICT.

In the left hippocampus model, both MICT (IRR=1.15, p<0.001, 95% CI [1.11, 1.20]) and C-HIIT (IRR=1.22, p<0.001, 95% CI [1.17, 1.26]) demonstrated higher rates than SC. C-HIIT outperformed MICT (IRR=1.06, p<0.001, 95% CI [1.03, 1.08]). When left hippocampus volume was considered as a baseline modulator, significant interactions emerged for MICT vs SC (IRR=0.94, p=0.001, 95% CI [0.90, 0.97]), C-HIIT vs SC (IRR=1.12, p<0.001, 95% CI [1.08, 1.17]), and C-HIIT vs MICT (IRR=1.20, p<0.001, 95% CI [1.17, 1.23]). This indicates that volume modulation reduced MICT benefits while enhancing C-HIIT benefits compared to SC, substantially increasing the C-HIIT advantage over MICT.

In the right hippocampus model, both MICT (IRR=1.15, p<0.001, 95% CI [1.11, 1.19]) and C-HIIT (IRR=1.20, p<0.001, 95% CI [1.16, 1.25]) showed higher rates than SC. C-HIIT outperformed MICT (IRR=1.05, p<0.001, 95% CI [1.03, 1.07]). When right hippocampus volume was considered as a baseline modulator, significant interactions emerged for MICT vs SC (IRR=0.90, p<0.001, 95% CI [0.87, 0.93]), C-HIIT vs SC (IRR=1.08, p<0.001, 95% CI [1.05, 1.11]), and C-HIIT vs MICT (IRR=1.20, p<0.001, 95% CI [1.17, 1.23]). This pattern mirrors the left hippocampus, with volume modulation reducing MICT benefits while maintaining C-HIIT benefits, substantially amplifying the C-HIIT advantage.

In the pars opercularis model, both MICT (IRR=1.27, p<0.001, 95% CI [1.23, 1.32]) and C-HIIT (IRR=1.29, p<0.001, 95% CI [1.25, 1.34]) achieved higher rates than SC. The MICT vs C-HIIT comparison was not significant. When pars opercularis volume was considered as a baseline modulator, the MICT vs SC interaction was not significant, while both C-HIIT vs SC (IRR=1.43, p<0.001, 95% CI [1.38, 1.49]) and C-HIIT vs MICT (IRR=1.39, p<0.001, 95% CI [1.35, 1.43]) showed significant interactions. This indicates that volume modulation specifically enhanced C-HIIT performance while leaving MICT effects unchanged, creating a significant C-HIIT advantage that was not present in the main effects.

In the brainstem model, both MICT (IRR=1.17, p<0.001, 95% CI [1.14, 1.21]) and C-HIIT (IRR=1.23, p<0.001, 95% CI [1.19, 1.27]) showed higher rates than SC. C-HIIT outperformed MICT (IRR=1.04, p<0.001, 95% CI [1.02, 1.07]). When brainstem volume was considered as a baseline modulator, significant interactions emerged for MICT vs SC (IRR=0.88, p<0.001, 95% CI [0.85, 0.91]), C-HIIT vs SC (IRR=1.27, p<0.001, 95% CI [1.23, 1.32]), and C-HIIT vs MICT (IRR=1.45, p<0.001, 95% CI [1.41, 1.48]). This indicates that volume modulation reduced MICT benefits while substantially enhancing C-HIIT benefits, creating the largest divergence between physical activity conditions with C-HIIT showing markedly superior performance.
